# Shisa7 phosphorylation regulates GABAergic transmission and neurodevelopmental behaviors

**DOI:** 10.1101/2021.08.02.454792

**Authors:** Kunwei Wu, Ryan David Shepard, David Castellano, Qingjun Tian, Lijin Dong, Wei Lu

## Abstract

GABA-A receptors (GABA_A_Rs) are crucial for development and regulation of the central nervous system. Altered GABAergic signaling is hypothesized to be involved in the pathophysiology of neurodevelopmental disorders. Nevertheless, how aberrant cellular and molecular mechanisms affect GABA_A_Rs in these diseases remain elusive. Recently, we identified Shisa7 as a GABA_A_R auxiliary subunit that modulates GABA_A_R trafficking, kinetics, and pharmacology, and discovered a phosphorylation site in Shisa7 (S405) critical for extrasynaptic α5-GABA_A_R trafficking and tonic inhibition. However, the role of S405 phosphorylation in the regulation of synaptic inhibition, plasticity, and behavior remains unknown. Here, we found that expression of a phospho-null mutant (Shisa7 S405A) in heterologous cells and neurons diminishes α2-GABA_A_R trafficking. Subsequently, we generate a Shisa7 S405A knock-in (KI) mouse line that displays reduced surface expression of GABA_A_Rs in hippocampal neurons. Importantly, both synaptic and tonic inhibition are decreased in KI mice. Moreover, chemically induced inhibitory long-term potentiation is impaired, highlighting a critical role of Shisa7 S405 in GABAergic plasticity. Lastly, KI mice exhibit enhanced locomotor activity and grooming associated with neurodevelopmental disorders. Collectively, our study reveals a phosphorylation site critical for Shisa7-dependent trafficking of synaptic and extrasynaptic GABA_A_Rs which contributes to behavioral endophenotypes displayed in neurodevelopmental disorders.

## Introduction

GABA-A receptors (GABA_A_Rs) are ligand-gated ion channels that are critical for regulating fast inhibitory neurotransmission within the central nervous system (CNS). Diversity in GABAergic inhibition arises from the pentameric assembly of 19 subunits into distinct receptor subtypes, which are broadly classified as synaptic or extrasynaptic GABA_A_Rs depending on their localization, as well as their biophysical and pharmacological properties (Olsen and Sieghart, 2009; Sieghart and Savić, 2018). Whereas synaptic GABA_A_Rs prominently contribute to phasic inhibition, extrasynaptic GABA_A_Rs have a more defined role in mediating tonic inhibition (Farrant and Nusser, 2005; Jacob et al., 2008). Due to their ubiquity across the CNS, it is not surprising that impaired GABA_A_R-mediated signaling is associated with a broad range of neurological and psychiatric disorders (Hines et al., 2012; B. Luscher et al., 2011; Möhler, 2006; Treiman, 2001). Therefore, defining mechanisms that regulate GABA_A_R activity and determining how perturbation of these processes are involved in pathophysiological conditions continue to be of high importance.

Historically, probing ligand-gated receptor function has been constrained to the level of their pore-forming subunits. However, this dogma has recently been challenged by the identification of protein complexes that coexist with native receptors. Accessory proteins and auxiliary subunits have been identified to interact with several voltage-gated ion channels and ligand-gated ion channels such as voltage-gated calcium channels, nicotinic acetylcholine receptors, and AMPA receptors (Maher et al., 2017). Adding to this growing list of receptor-interacting proteins, GABA_A_R-associated transmembrane proteins have become the most recent addition and consist of lipoma HMGIC fusion partner-like 4 (LH4), Cleft lip and palate transmembrane protein 1 (Clptm1), and Shisa7 (Castellano et al., 2020; Han et al., 2020). Shisa7 is a single-pass transmembrane protein that was identified in a proteomic screen and as part of native GABA_A_R complexes (Nakamura et al., 2016). Indeed, our initial study identified that Shisa7 localizes at inhibitory synapses and directly interacts with several GABA_A_R subunits (Han et al., 2019). Coexpression of Shisa7 with specific GABA_A_R subtypes in heterologous cells resulted in enhanced GABA-evoked, whole-cell currents. Moreover, loss of Shisa7 impaired phasic GABAergic transmission in hippocampal CA1 pyramidal neurons. Interestingly, Shisa7 also accelerated IPSC decay kinetics and loss of Shisa7 blunted the pharmacological and behavioral effects of diazepam, a prototypical benzodiazepine. In addition, we found that Shisa7 regulated trafficking of extrasynaptic α5-GABA_A_Rs, but not δ-GABA_A_Rs (Wu et al., 2021b). Specifically, loss of Shisa7 resulted in diminished trafficking of α5-GABA_A_Rs and tonic currents. Interestingly, S405 on Shisa7 was identified as a critical phosphorylation site for protein kinase A (PKA) and PKA-mediated phosphorylation of S405 promoted Shisa7-dependent forward trafficking of α5-GABA_A_Rs.

Recently, accumulating evidence has shown that disrupted coordination of excitatory and inhibitory inputs throughout the CNS are associated with neurodevelopmental disorders such as autism (ASD), attention-deficit hyperactivity disorder (ADHD), and obsessive-compulsive disorder (OCD) (Gatto and Broadie, 2010; Lee et al., 2017). Although a broad range of differential mechanisms link altered CNS function with neurodevelopmental disorders, there are still shared characteristics that implicate overlapping similarities in behavioral presentation such as impairments in social interaction, deficits in language/communication skills, and repetitive or altered vigilance with attention-based tasks (Coghlan et al., 2012) which can impede proper diagnosis and treatment (Brem et al., 2014; Leyfer et al., 2006). Additionally, rodent models of ASD/ADHD/OCD share common behavioral endophenotypes such as impulsivity, hyperactivity, impaired social interaction, deficits in learning and memory, and repetition of behaviors (Ahmari, 2016; Kazdoba et al., 2016). Although polygenic inheritance likely drives the etiology and core behavioral traits observed in ASD/ADHD/OCD, studies implicate alterations in GABA_A_R expression. In particular, changes in the 15q11-13 chromosomal region have been linked to ASD which contains the *GABRB3, GABRA5*, and *GABRG3* genes that encode for β3, α5, and γ3 subunits, respectively, (Tager-Flusberg et al., 2001). Additionally, alterations in the *GABRB3* gene are associated with ASD (Buxbaum et al., 2002). Moreover, it has been reported that β3 expression (Samaco et al., 2005) and mRNA for the various GABA_A_R subunits are decreased in autistic individuals, albeit in discrete brain regions (Fatemi et al., 2014, 2009). Rodent knock-out (KO) studies also suggest that GABA_A_R loss could be involved in neurodevelopmental disorders. For example, KO of α5-GABA_A_Rs results in ASD-like behaviors in mice, indicating a critical role for tonic inhibition (Mesbah-Oskui et al., 2017; Zurek et al., 2016). In line with clinical observations, β3 KO mice also display some endophenotypes such as reduced sociability, hyperactivity, seizure susceptibility, and deficits in attention, learning and memory (Culiat et al., 1994; DeLorey et al., 2008; Homanics et al., 1997). Lastly, alterations in grooming behavior are seen in a wide variety of ASD models (Gandhi and Lee, 2021) which can be bi-directionally regulated using GABA_A_R pharmacology (Kalueff et al., 2016). Taken together, these observations suggest that altered GABA_A_R function might be involved in neurodevelopmental disorders and require further investigation.

Based on our finding that S405 phosphorylation is important for Shisa7-dependent trafficking of α5-GABA_A_Rs (Wu et al., 2021b), we investigated the role of this residue in synaptic inhibition, inhibitory synaptic plasticity, and behavior. We found that in both heterologous cells and hippocampal neuronal cultures expression of Shisa7 S405A mutant decreased surface α2-GABA_A_Rs. Following these findings, we generated a knock-in (KI) mouse line encoding the Shisa7 phospho-null single-point mutation (S405A) (hereafter KI mice). Interestingly, KI mice displayed a reduction in surface expression of α2-GABA_A_Rs in hippocampal neurons along with impaired phasic inhibition compared to WT. Significantly, chemically-induced inhibitory long-term potentiation (chem-iLTP) was blunted in hippocampal CA1 neurons in KI mice. Lastly, we behaviorally phenotyped KI mice and observed locomotor hyperactivity and increased grooming behavior. Taken together, our data demonstrate that a specific residue, S405, in Shisa7 C-terminus plays an unappreciated, but critical role in regulating synaptic inhibition and plasticity, and impairment of Shisa7 S405-dependent regulation of tonic and phasic inhibition *in vivo* may contribute to specific behavioral endophenotypes shared among ASD/ADHD/OCD.

## Results

### Expression of Shisa7 S405A mutant reduces surface expression of α2-GABA_A_Rs

Given that Shisa7 promoted trafficking of synaptic GABA_A_Rs (Han et al., 2019), we investigated whether Shisa7 S405, a site previously found to be a phosphorylation substrate of PKA and important for trafficking of extrasynaptic α5-GABA_A_Rs (Wu et al., 2021b), is also critical for synaptic GABA_A_R trafficking. Using immunocytochemistry, we observed that heterologous cells expressing α2β3γ2 GABA_A_Rs had increased surface α2 expression when co-transfected with Shisa7, but not with the phospho-null S405A mutant (**Figure 1A**), indicating that the regulation of α2-GABA_A_R expression at the surface by Shisa7 is dependent on S405 phosphorylation. Consistently, co-expression of Shisa7 S405A in heterologous cells transfected with α2β3γ2 decreased GABA-evoked whole-cell currents (**Figure 1B**). We next examined whether Shisa7 S405 modulated α2-GABA_A_R abundance at the surface in neurons. We found that expression of the Shisa7 S405A mutant significantly reduced surface α2-GABA_A_Rs in cultured hippocampal neurons (**Figure 1C**). Taken together, these data extend our previous observations that Shisa7 promotes GABA_A_R trafficking to the cell surface (Han et al., 2019; Wu et al., 2021b) and show that Shisa7 S405 regulates trafficking of both synaptic and extrasynaptic GABA_A_Rs.

**Figure 1.**
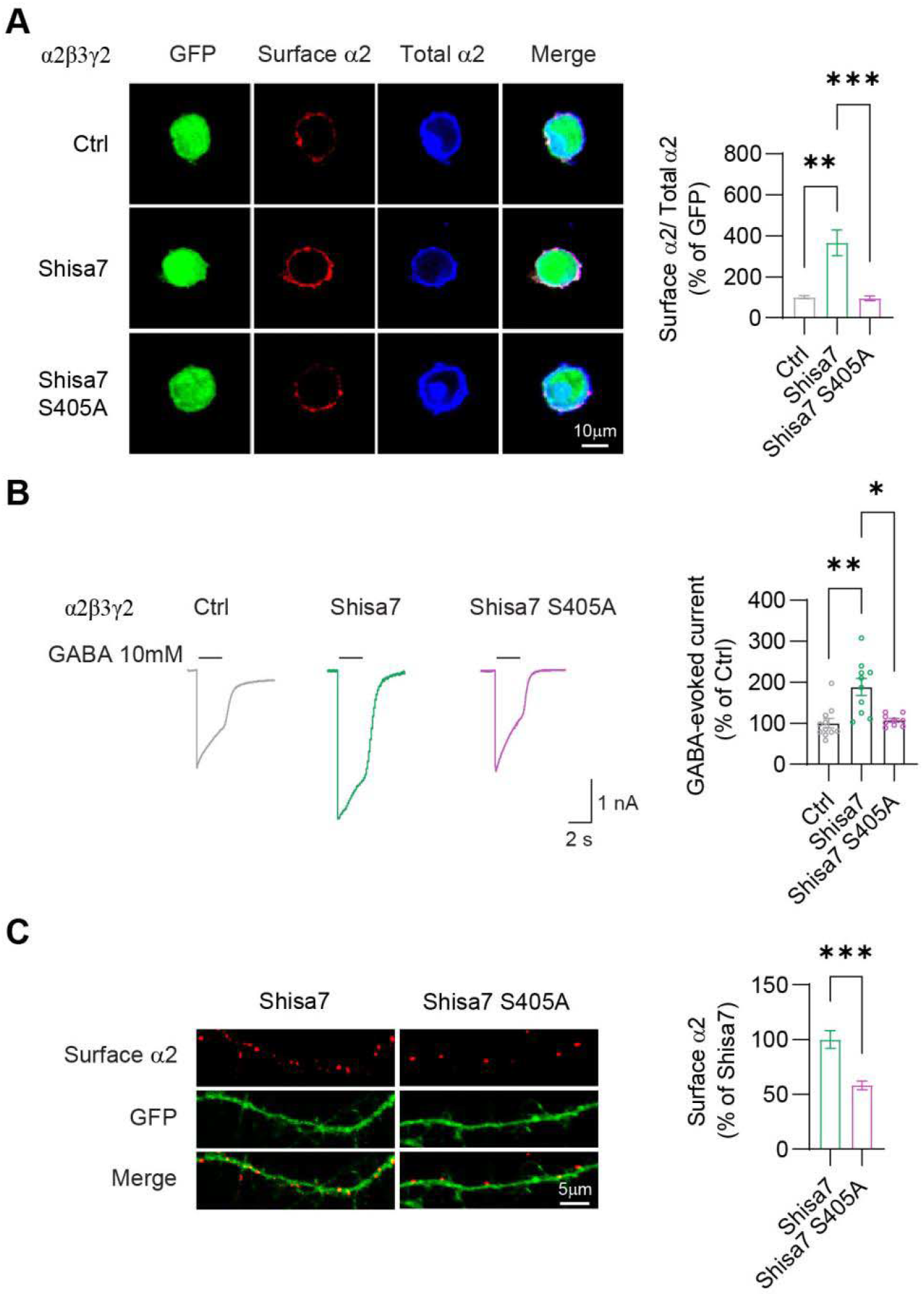
Expression of Shisa7 S405A mutant reduces cell surface expression of α2-GABA_A_Rs. **A**. Surface α2 expression is increased following co-transfection of Shisa7, but not with S405A mutant in HEK293T cells expressing α2β3γ2 (Ctrl, n = 18; Shisa7, n = 20; Shisa7 S405A, n = 22, Kruskal-Wallis test with Dunnett’s multiple comparison test; Ctrl versus Shisa7, p = 0.0046; Shisa7 versus Shisa7 S405A, p = 0.0002). **B**. S405A significantly reduces GABA-evoked whole-cell currents compared to Shisa7 in HEK293T cells expressing α2β3γ2 (Ctrl, n = 11; Shisa7, n = 10; Shisa7 S405A, n = 9, Kruskal-Wallis test with Dunnett’s multiple comparison test; Ctrl versus Shisa7, p = 0.0011; Shisa7 versus Shisa7 S405A, p = 0.041). **C**. Expression of S405A mutant decreases surface α2 compared to those expressed with Shisa7 in hippocampal neurons (Shisa7, n = 20; Shisa7 S405A, n = 26, Mann-Whitney U test; p = 0.0003). Error bars indicate S.E.M. *p < 0.05; **p < 0.01; ***p < 0.001.

### Shisa7 S405 is required for surface expression of GABA_A_Rs and GABAergic transmission in hippocampal neurons

To further corroborate the role of Shisa7 S405 in the regulation of GABA_A_R trafficking and GABAergic transmission, we generated a Shisa7 S405A KI mouse line that is phospho-deficient at S405 (**Supplementary Figures 1A, B**). To examine whether the mutation altered the overall expression of Shisa7 in the hippocampus, we first conducted Western blots using hippocampal lysates which demonstrated that Shisa7 expression was not significantly changed in our mouse line (**Supplementary Figure 1C**). To investigate the impact of the S405A mutation on GABA_A_R surface expression, we performed immunocytochemical experiments in hippocampal neuronal cultures. We found that α2 surface expression was significantly reduced in KI neurons (**Figure 2A**), indicating that phosphorylation of S405 is critical for surface expression of a major synaptic GABA_A_R subtype. Similarly, surface expression of extrasynaptic α5-GABA_A_Rs was also significantly decreased (**Figure 2B**), suggesting that Shisa7 S405 is involved in trafficking of both synaptic and extrasynaptic GABA_A_Rs. We further confirmed this observation by utilizing electrophysiological recordings which showed decreased GABA-evoked whole-cell currents in dissociated hippocampal cultures prepared from KI mice (**Figure 2C**).

**Figure 2.**
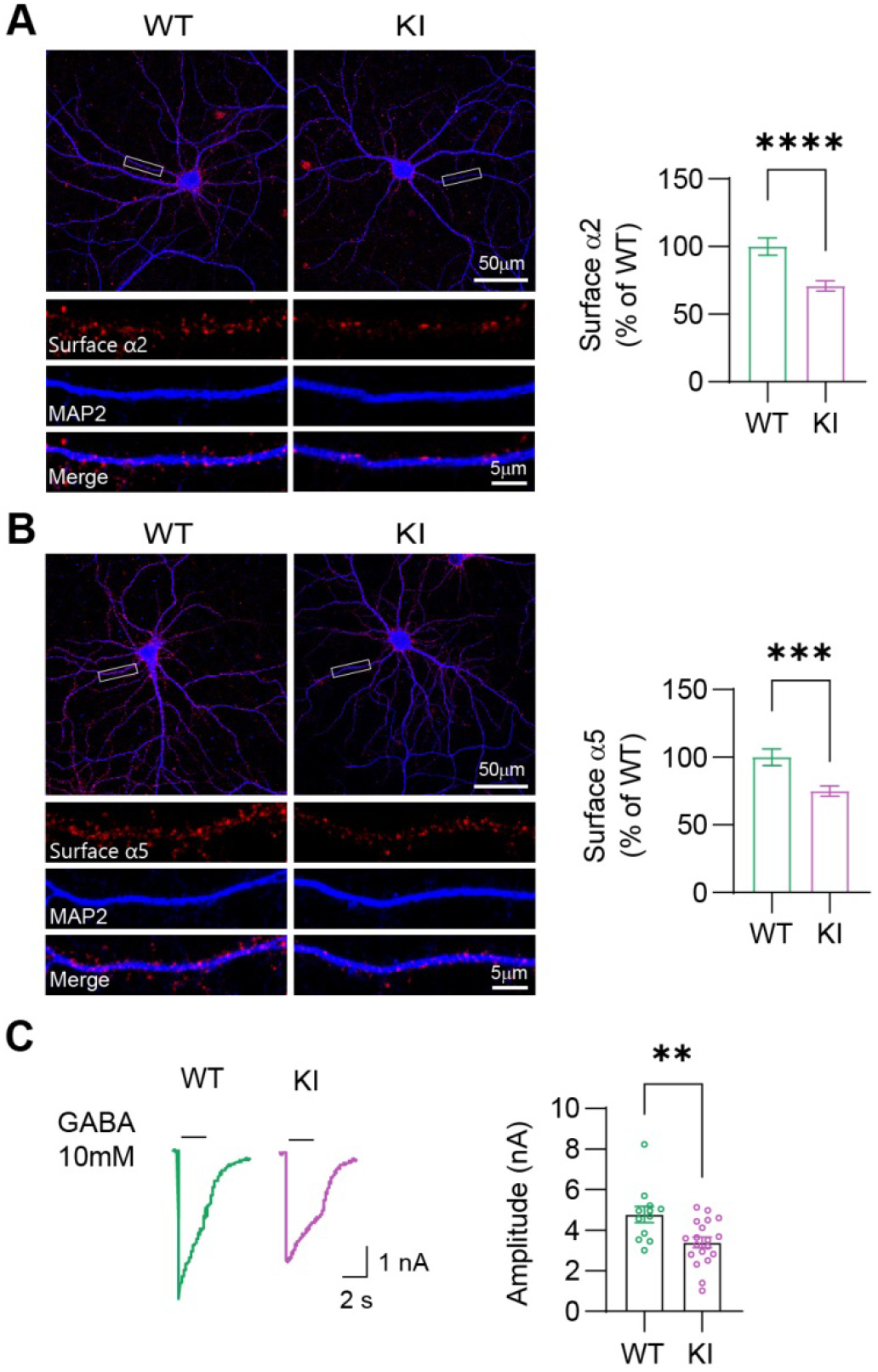
Shisa7 S405A KI mice exhibit decreased cell surface expression of GABA_A_Rs. **A**. Surface α2 expression in cultured hippocampal neurons from KI mice is reduced (WT, n = 31; KI, n = 31, Mann-Whitney U test; p < 0.0001) **B**. Surface α5 expression in cultured hippocampal neurons from KI mice is reduced (WT, n = 27; KI, n = 32, Mann-Whitney U test; p = 0.0004) **C**. GABA-evoked whole-cell currents in cultured hippocampal neurons from KI mice are decreased (WT, n = 12; KI, n = 19, t test; p = 0.0049). Error bars indicate S.E.M. **p < 0.01; ***p < 0.001; ****p < 0.0001.

Next, we evaluated whether both GABA_A_R-mediated synaptic and tonic currents were altered in dissociated hippocampal cultures prepared from KI mice. Although no significant alterations in mIPSC amplitude were observed, there was a strong decrease in mIPSC frequency, as well as diminished tonic currents in KI hippocampal neurons (**Figures 3A-D**). We further investigated whether mIPSCs were affected in CA1 pyramidal neurons in acute hippocampal slices prepared from KI mice. Congruent with the evidence from dissociated hippocampal cultures, we observed a reduction in mIPSC frequency, but not amplitude, in KI mice (**Figures 3E-G**). Therefore, S405 phosphorylation is required for Shisa7-dependent trafficking of synaptic α2- and extrasynaptic α5-GABA_A_Rs in hippocampal neurons.

**Figure 3.**
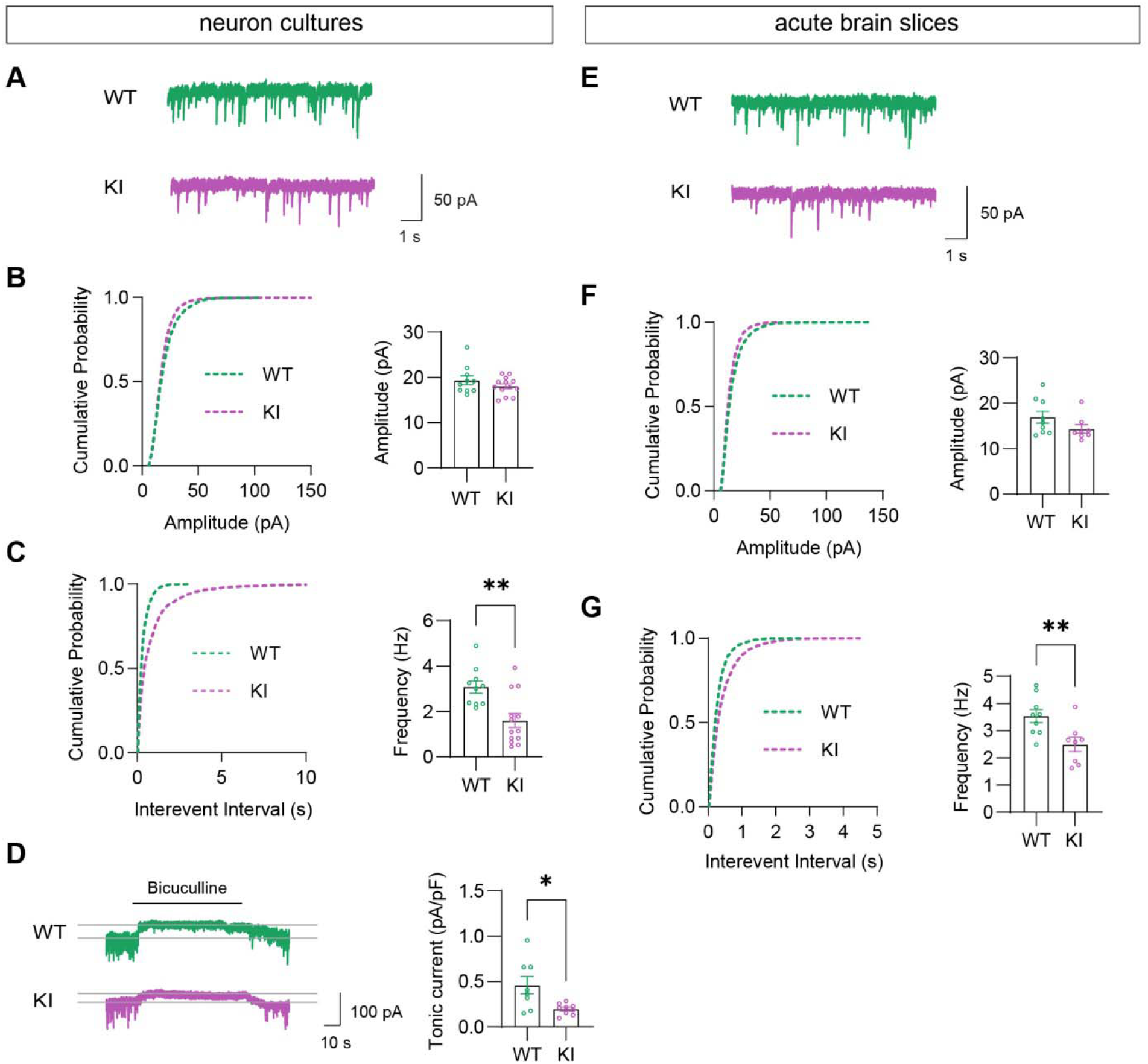
Shisa7 S405A KI mice exhibit impaired phasic and tonic inhibition. **A**. Representative mIPSC traces from neuronal cultures. **B. and C**. mIPSC frequency is diminished in KI mice with no change in mIPSC amplitude (WT, n = 10; KI, n = 13, Mann-Whitney U test; p = 0.0024) **D**. Tonic inhibition is reduced in neuronal cultures from KI mice (WT, n = 8; KI, n = 9, t test; p = 0.0129) **E**. Representative mIPSC traces from CA1 neurons in acute hippocampal slices. **F-H**. mIPSC frequency is decreased onto CA1 neurons in KI mice with no change in mIPSC amplitude (WT, n = 9; KI, n = 8, Mann-Whitney U test; p = 0.0099). Error bars indicate S.E.M. *p <0.05; **p <0.01.

### iLTP is impaired in KI mice

Regulation of GABAergic plasticity is dependent on the surface abundance of GABA_A_Rs and thus trafficking processes can affect the expression of iLTP (Castillo et al., 2011; Bernhard Luscher et al., 2011). To this end, we adopted a strategy (Marsden et al., 2007) (chem-iLTP) to assess whether iLTP was altered in hippocampal neurons from KI mice. Specifically, following transient application of NMDA (3 min, 20 μM) in the bath perfusate, we examined mIPSC amplitude in a time-dependent manner up to 30 min after NMDA exposure in CA1 pyramidal neurons from acute hippocampal slices. In agreement with previous studies (Marsden et al., 2007; Petrini et al., 2014; Wiera et al., 2021), brief NMDA exposure to hippocampal slices was sufficient to induce a persistent increase in mIPSC amplitude over a 30-min period in WT mice (**Figure 4A**). Strikingly, mIPSC amplitude in CA1 pyramidal neurons in KI mice 30 min post-NMDA application did not significantly increase past baseline levels (**Figures 4B, C**). These findings suggest that Shisa7 S405 phosphorylation is a requirement for activity-dependent trafficking of GABA_A_Rs and iLTP in CA1 pyramidal neurons.

**Figure 4.**
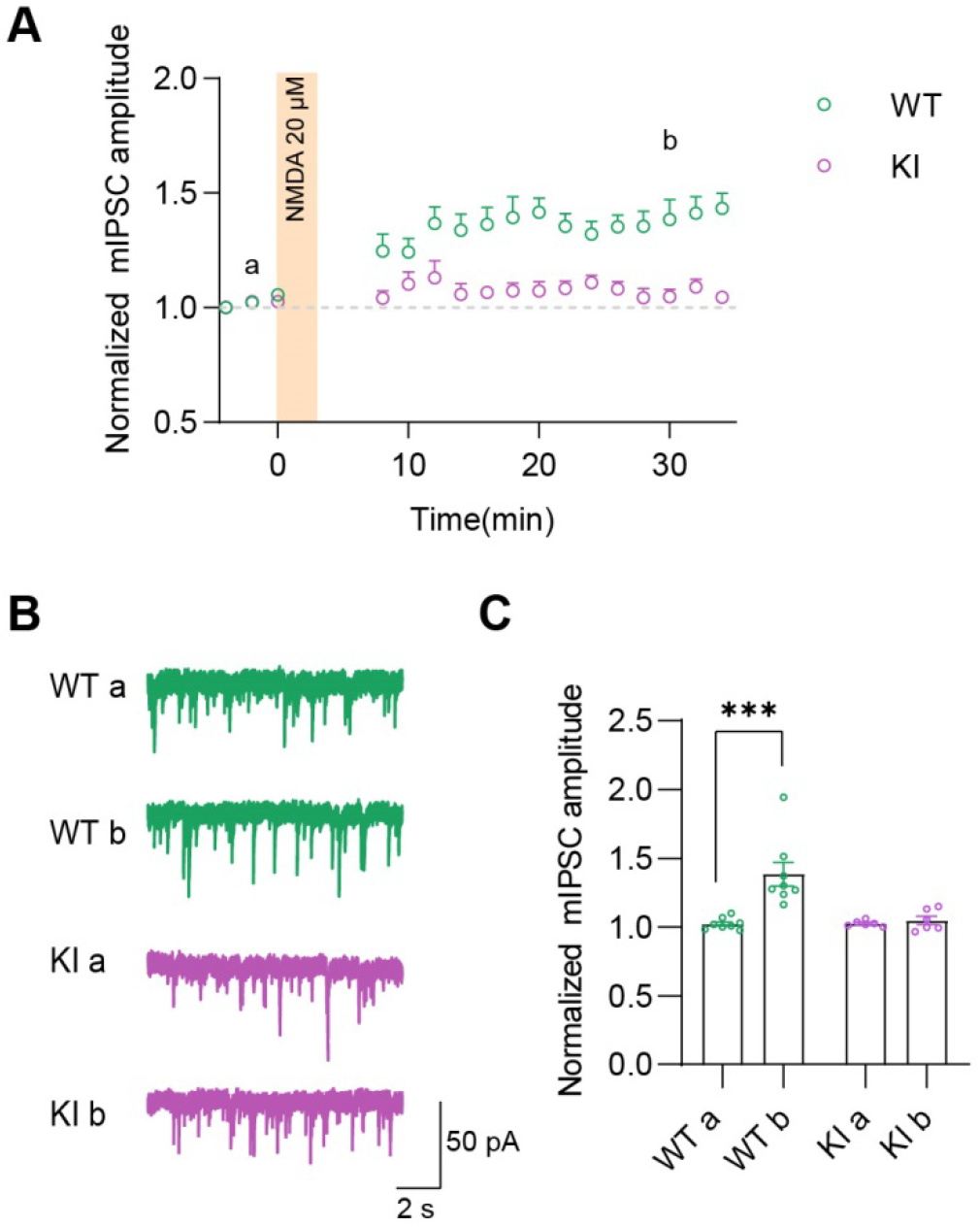
iLTP is impaired in CA1 neurons from Shisa7 S405A KI mice. **A**. Bath application of NMDA (3 min, 20 µM) increases mIPSC amplitude for up to 30 min compared to baseline in WT mice, but not KI mice. **B**. Representative traces of mIPSCs from CA1 neurons in acute hippocampal slices at baseline and then 30 min following bath-application of NMDA. **C**. Bath application of NMDA (3 min, 20 μM) increases mIPSC amplitude in WT mice only (WT, n = 8; KI, n = 6, two-way ANOVA, F _16, 204_ = 2.324, p = 0.0036 with Sidak’s multiple comparison test; WT a versus WT b, p = 0.0005;). Error bars indicate S.E.M. ***p <0.001.

### KI mice display locomotor hyperactivity and increased grooming behavioral endophenotypes

Given that KI mice displayed impaired inhibitory transmission and iLTP, we examined the consequences of Shisa7 S405A mutation on animal behaviors. We first investigated whether KI mice had any deficits in learning and memory using two behavioral paradigms. In order to assess short-term spatial working memory, we utilized the Y-maze. Mice were allowed to freely explore for 5 min and spontaneous alternation behavior was defined as consecutive entry into all three arms without repetition (Hughes, 2004). We observed that KI mice exhibited no impairment of spontaneous alternation behavior compared to WT mice (**Figure 5A**). Next we assessed object recognition memory using the Novel Object Recognition (NOR) test. During the test session, WT and KI mice performed similarly, spending more time with the novel object compared to the familiar object (**Figure 5B**).

**Figure 5.**
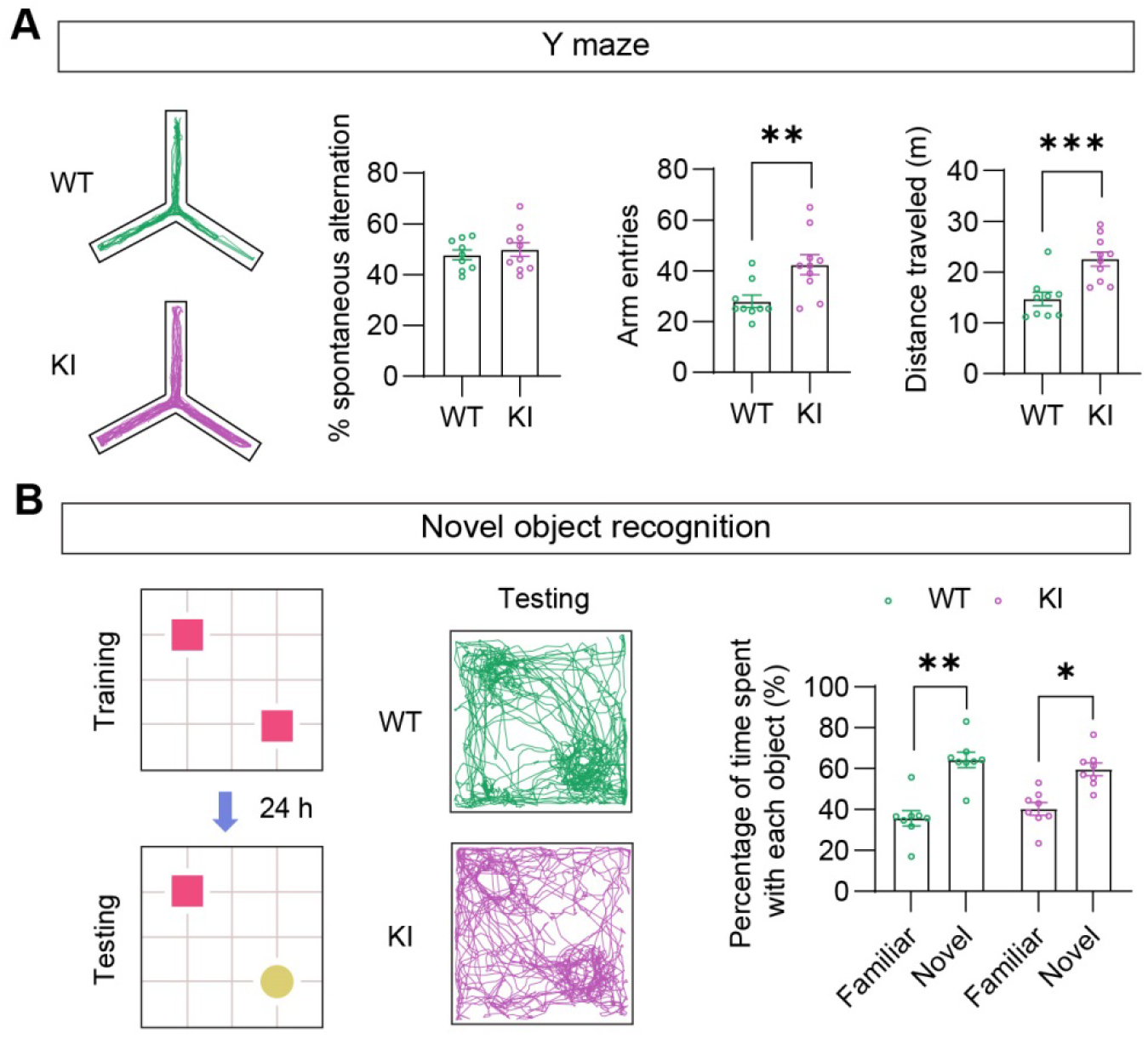
Short-term spatial memory and object recognition memory are intact in KI mice. KI mice have increased number of arm entries (WT, n = 9; KI, n=10, t test; p = 0.0077) and total distance traveled (WT, n = 9; KI, n=10, t test; p = 0.0009), but no change in spontaneous alternation among the arms in the Y maze (WT, n = 9; KI, n=10, t test) **B**. KI mice do not show signs of impaired learning and memory in the NOR test (WT, n = 8; KI, n=8, two-way ANOVA, F _1, 14_ = 24.34, p = 0.0002 with Sidak’s multiple comparison test; WT: Familiar versus WT: Novel, p = 0.0019; KI: Familiar versus KI: Novel, p = 0.0277). Error bars indicate S.E.M. *p < 0.05; **p < 0.01; ***p < 0.001.

From these two assays, KI mice did not display any noticeable deficits in learning and memory. However, we observed that KI mice had increased arm entries and increased total distance traveled during the Y-maze test (**Figure 5A**), suggesting that KI mice displayed locomotor hyperactivity. To confirm this, we measured general locomotor activity using the Open Field Test (OFT) and homecage activity recordings. During the 30-min OPT, the total distance traveled by KI mice was significantly larger than WT mice (**Figure 6A**) In the 24-h homecage activity recordings, KI mice also traveled greater distances as indicated by increased beam interruptions compared to WT mice; although, the increased movement seemed to be more prevalent during “lights off” (ZT12-24; **Figure 6B**). Overall, these behavioral data indicate that KI mice have a hyperactive locomotor phenotype. Since hyperactivity has been observed in animal models of ASD/ADHD/OCD (Ahmari, 2016; Kazdoba et al., 2016), we proceeded to further characterize KI mice by assessing repetitive and compulsive behaviors using Nestlet Shredding Test (NST), grooming time, and Marble Burying Test (MBT) (Angoa-Pérez et al., 2013; Kalueff et al., 2016). Although KI mice performed similarly to WT mice in the NST and MBT, KI mice spent a greater amount of time grooming during a 10-min period (**Figures 6C-F**). Given the co-morbidity of depression (Magnuson and Constantino, 2011) and sleep disturbances (Malow and McGrew, 2008) with ASD/ADHD/OCD, we also assessed pro-depressive coping styles (learned helplessness) using the Forced Swim Test (FST) and Piezoelectric sleep recordings to monitor sleep, respectively. Both the FST and sleep recordings yielded no difference between WT and KI mice with respect to immobility time and total sleep time, respectively (**Figures 6E, G**). In sum, impaired Shisa7-dependent trafficking of GABA_A_Rs and iLTP in KI mice with the phospho-null mutation S405A produced endophenotypes of neurodevelopmental disorders, specifically locomotor hyperactivity and increased grooming behavior.

**Figure 6.**
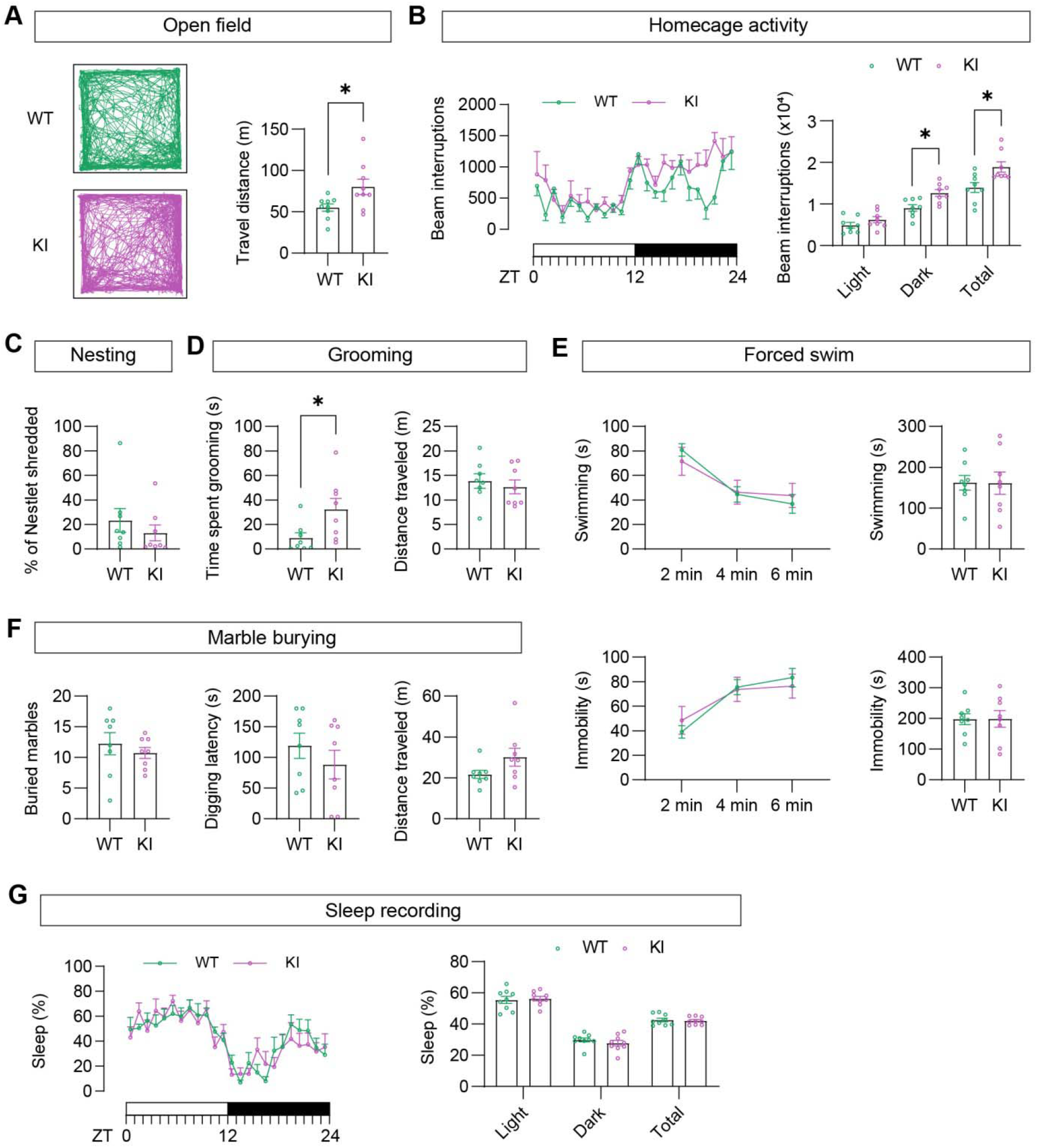
Shisa7 S405A KI mice display locomotor hyperactivity and increased grooming behavioral endophenotypes. **A**. KI mice have increased locomotion compared to WT mice in OFT (WT, n = 9; KI, n = 9, t test; p = 0.0265) **B**. KI mice display more activity (# of beam interruptions) in the homecage driven by enhanced activity during “lights off” (ZT12-24); WT, n = 8; KI, n = 8, two-way ANOVA, F _2, 28_ = 5.329, p = 0.0109 with Sidak’s multiple comparison test; Dark: WT versus Dark: KI, p = 0.0191; Total: WT versus Total: KI, p = 0.0392) **C**. There is no difference in the amount of nestlet shredded between WT and KI mice (WT, n = 8; KI, n = 8, Mann-Whitney U test) **D**. KI mice have increased grooming behavior compared to WT (WT, n = 8; KI, n = 8, Mann-Whitney U test; p = 0.0103) **E**. There is no difference in burying activity between WT and KI mice (WT, n = 8; KI, n = 8, t test) **F**. KI mice do not display learned helplessness behavior in comparison to WT mice (WT, n = 8; KI, n = 8, t test) **G**. Sleep behavior is not different between WT and KI mice (WT, n = 9; KI, n = 9, two-way ANOVA with Sidak’s multiple comparison test). Error bars indicate S.E.M. *p<0.05.

## Discussion

GABA_A_Rs are critical for the regulation of neuronal excitability across the CNS and deficits in GABA_A_R-mediated signaling are associated with a myriad of psychiatric conditions (B. Luscher et al., 2011; Möhler, 2006; Treiman, 2001). Therefore, it remains essential to understand the mechanistic processes by which GABA_A_Rs are trafficked to the synaptic and extrasynaptic compartments. In this study, we have expanded upon our previous observations regarding the role of Shisa7 in GABA_A_R trafficking (Han et al., 2019; Wu et al., 2021b) by providing specific insight into the role for S405 phosphorylation. We have demonstrated that S405 phosphorylation is not only instrumental for trafficking of extrasynaptic GABA_A_Rs (Wu et al., 2021b), but is also important for trafficking of synaptic GABA_A_Rs. Importantly, we observed that GABA_A_R trafficking deficits in Shisa7 S405A KI mice not only resulted in diminished GABAergic transmission, but also blunted NMDAR-mediated iLTP. Lastly, we identified specific endophenotypes (locomotor hyperactivity and increased grooming) in Shisa7 KI mice which are associated with ASD/ADHD/OCD. Collectively, this study extends our knowledge regarding the role of Shisa7-dependent regulation of both synaptic and extrasynaptic GABA_A_Rs and reveals that behavioral consequences of not allowing Shisa7 S405 to be phosphorylated are locomotor hyperactivity and longer grooming periods.

### S405 phosphorylation is required for Shisa7-Dependent Trafficking of α2-GABA_A_Rs

Trafficking of GABA_A_Rs to inhibitory synapses is mediated by various molecular and cellular processes which include a number of binding partners such as GABA_A_R associated protein (GABARAP), clathrin adaptor protein 2 (AP2) complex, *N*-ethylmaleimide-sensitive factor (NSF), Phospholipase C-related catalytically inactive proteins (PRIPs) and radixin (Jacob, 2019; Jacob et al., 2008; Bernhard Luscher et al., 2011). The most recent advancements to our knowledge pertaining to GABA_A_R trafficking have been the role of novel GABA_A_R-associated transmembrane proteins as critical regulators (Castellano et al., 2020; Han et al., 2020). For example, LH4 KO mice show decreased GABA_A_R synaptic clustering (Davenport et al., 2017; Yamasaki et al., 2017). In contrast, Clptm1 KO mice show increased GABA_A_R surface expression due to the role of endogenous Clptm1 as a negative regulator of GABA_A_R forward trafficking (Ge et al., 2018). Interestingly, LH4 showed specificity for γ2 -GABA_A_Rs by forming a tripartite complex with neuroligin-2, whereas Clptm1 associated with both synaptic and extrasynaptic GABA_A_Rs (Yamasaki et al., 2017). However, the critical residues required for these transmembrane proteins to exert their effects on GABA_A_R trafficking have not been reported. Additionally, it is not known whether posttranslational modifications (PTMs) also affect their trafficking ability. Our initial study showing that Shisa7 was an auxiliary subunit also identified the GABA_A_R interacting domain (GRID) as a critical region on Shisa7 required for synaptic GABA_A_R trafficking (Han et al., 2019). We subsequently showed that Shisa7 regulated tonic inhibitory currents in CA1 pyramidal neurons via promoting α5-GABA_A_R exocytosis (Wu et al., 2021b). Importantly, Shisa7 regulation of tonic inhibition and α5-GABA_A_R trafficking requires PKA-dependent phosphorylation of S405. Here, we report that S405 phosphorylation is also needed for Shisa7-dependent trafficking of synaptic α2-GABA_A_Rs. While this demonstrates a crucial residue for Shisa7-dependent regulation of both synaptic and extrasynaptic GABA_A_Rs, it also suggests that PTMs, specifically phosphorylation, on Shisa7 are critical for how Shisa7 impacts GABA_A_Rs. Further investigation into additional key residues on Shisa7 and how other types of PTMs influence Shisa7-dependent regulation of GABA_A_Rs are warranted.

### S405 KI mice have impaired phasic and tonic inhibition

Mechanistically, GABA_A_Rs control neuronal excitability due to their high chloride permeability which influxes and hyperpolarizes neurons upon activation. However, GABAergic signaling is diverse and is largely dependent on subunit composition and receptor localization (Olsen and Sieghart, 2009; Sieghart and Savić, 2018). In addition, the strength of GABAergic inhibition is directly related to GABA_A_R abundance at the cell surface. In the case of GABA_A_R-associated transmembrane proteins, they all have been shown to play a critical role in inhibitory neurotransmission (Castellano et al., 2020; Han et al., 2020). In LH4 KO mice, recordings from hippocampal CA1 pyramidal neurons show decreased mIPSCs, indicating that LH4 is poised to directly affect the strength of synaptic inhibition (Davenport et al., 2017; Yamasaki et al., 2017). Given that Clptm1 restricts forward trafficking of both synaptic and extrasynaptic GABA_A_Rs, overexpression and knockdown of Clptm1 diminished and enhanced both mIPSCs and tonic currents, respectively (Ge et al., 2018). We previously showed that KO of Shisa7 can diminish both phasic (Han et al., 2019) and tonic inhibition (Wu et al., 2021b) in hippocampal neurons. Since we identified S405 phosphorylation as a regulator of tonic inhibition (Wu et al., 2021b), we explored whether phosphorylation of this residue also impacted phasic inhibition in hippocampal neurons. Using a newly generated KI mouse line, we observed a reduction in both phasic and tonic inhibition in hippocampal neurons. These decreases in GABAergic inhibition are attributed to diminished surface expression of α2- and α5-GABA_A_Rs. These data recapitulate our previous findings that Shisa7 S405 is indeed critical for tonic currents and provide further evidence that prohibiting S405 phosphorylation consequently results in diminished mIPSCs. Therefore, these data suggest that Shisa7 S405 phosphorylation has a comprehensive role in controlling GABAergic inhibition that encompasses both phasic and tonic inhibition.

### iLTP is impaired in Shisa7 S405A KI mice

Synaptic plasticity has been proposed to be the cellular basis of learning and memory that involves diverse processes that lead to either strengthening or weakening of synapses. Although the molecular mechanisms underlying the regulation of excitatory synaptic plasticity have been extensively studied (Diering and Huganir, 2018; Huganir and Nicoll, 2013; Maffei, 2018), much less is known about the regulation of inhibitory synaptic plasticity. Accumulating studies have revealed a critical role for NMDARs in the regulation of inhibitory synapse development and function (Gu et al., 2016; Gu and Lu, 2018; Horn and Nicoll, 2018; Wu et al., 2021a), and the induction of iLTP (Chiu et al., 2018; Marsden et al., 2007; Petrini et al., 2014; Wiera et al., 2021). It has been shown that NMDAR-mediated iLTP requires synaptic recruitment of gephyrin, CaMKII activation, and synaptic insertion of β2/3-containing GABA_A_Rs (Chiu et al., 2018; Marsden et al., 2007; Petrini et al., 2014). Recent studies have also identified a number of transmembrane accessory proteins for GABA_A_Rs (Han et al., 2020). However, the role of these transmembrane proteins in iLTP remained unknown. We have now demonstrated a critical role of Shisa7 in the regulation of iLTP. Specifically, in Shisa7 S405A KI mice, iLTP in hippocampal CA1 pyramidal neurons was blunted, indicating that synaptic recruitment of GABA_A_Rs during iLTP depends on a Shisa7 S405 phosphorylation process. Since Shisa7 S405 is the substrate of PKA, our data also suggest a potential role of PKA in iLTP. Currently, how Shisa7 S405 dependent pathway functionally interacts with gephyrin and CaMKII dependent processes to regulate iLTP remains unknown. Additionally, it is worth noting that our iLTP experiment was measured through recording mIPSCs from CA1 pyramidal neurons. Therefore, it remains to be determined whether Shisa7 can regulate iLTP in a synapse-specific manner. It has recently been reported that the expression of iLTP in hippocampal and cortical pyramidal neurons is input-dependent (Chiu et al., 2018; Udakis et al., 2020). It will be important to understand how Shisa7 regulates GABAergic plasticity in a synapse-specific context.

In addition to iLTP, other forms of plasticity including iLTD and homeostatic plasticity have been identified at inhibitory synapses (Castillo et al., 2011; Chiu et al., 2019; Hartman et al., 2006; Kilman and Van Rossum, 2002). Thus, it will be valuable to investigate iLTD and inhibitory homeostatic plasticity within the context of Shisa7 regulation. Notably, our recent study identified a specific role for Shisa7 and Shisa7 S405 in regulating homeostatic plasticity of tonic inhibitory currents in hippocampal neurons (Wu et al., 2021b). It would be interesting to examine whether inhibitory synaptic scaling is also dependent on Shisa7 S405 phosphorylation.

### KI mice display locomotor hyperactivity and increased grooming behavioral endophenotypes

GABA_A_Rs are critical in the regulation of neural circuit function of the CNS (Braat and Kooy, 2015; Brickley and Mody, 2012; Hines et al., 2012). Given our findings of impaired inhibitory transmission and iLTP in KI mice reported here, we performed a battery of behavioral tests to examine the consequences of Shisa7 S405A mutation on animal behaviors. We first investigated whether learning and memory behaviors were altered in KI mice. Interestingly, although KI mice did not display any deficits in both the Y-maze and NOR tests, these mice exhibited increased locomotor activity in the Y-maze. We further confirmed this increased locomotor activity using the OFT and by examining homecage activity. Given that hyperactivity has been associated with ASD/OCD/ADHD (Ahmari, 2016; Kazdoba et al., 2016), we utilized behavioral paradigms to determine whether our KI mouse line had any additional endophenotypes associated with these neurodevelopmental disorders. Although we observed no difference in nestlet shredding or marble burying, two common assays for repetitive animal behaviors (Angoa-Pérez et al., 2013), we did observe a greater propensity to groom in KI mice compared to WT. This intriguing finding is in line with pharmacological experiments looking at grooming behavior in that antagonism of GABA_A_Rs has been shown to precipitate greater grooming behavior (Kalueff et al., 2016). Therefore, deficits in Shisa7-dependent trafficking of GABA_A_Rs could be impacting discrete brain regions whereby GABA_A_Rs influence grooming behaviors. Additionally, clinical evidence has suggested that individuals afflicted with ASD/OCD/ADHD are more likely to develop mood disorders, such as depression (Magnuson and Constantino, 2011). Indeed, animal models of ASD/OCD/ADHD have been shown to recapitulate depressive-like behaviors, such as increased immobility in the FST (Commons et al., 2017). However, we did not see any differences in immobility time from KI mice compared to WT, demonstrating that these animals do not display any pro-depressive behaviors. Lastly, because sleep disturbances are often comorbid with many neurodevelopmental disorders and due to our previous observations that tonic inhibitory currents in hippocampal neurons differs based on sleep-wake states (Wu et al., 2021b), we monitored sleep behavior using a piezoelectric monitoring system (Mang et al., 2014; Yaghouby et al., 2016). In contrast to our previous study which showed decreased sleep in Shisa7 KO mice (Wu et al., 2021b), we did not observe this effect here. One major difference between our two studies is the former had no Shisa7 present throughout the development whereas the current study here has Shisa7, but with the S405 site rendered incapable of being phosphorylated. Although we note that Shisa7 S405 is important for trafficking of synaptic and extrasynaptic GABA_A_Rs, it is possible that S405-independent mechanisms are involved in Shisa7 regulation of sleep behavior. Additionally, since Shisa7 regulates trafficking of both synaptic and extrasynaptic GABAARs, it is difficult to conclude whether these endophenotypes are the result of impaired Shisa7-dependent trafficking of synaptic or extrasynaptic GABA_A_Rs. For example, enhanced grooming time has been observed in α5-GABA_A_R KO mice (Zurek et al., 2016), but loss of α5-GABA_A_Rs has also been observed to enhance learning (Collinson et al., 2002). Currently, the mechanisms underlying the role of Shisa7 S405 in animal behavior remain unknown. It is possible that cell-type specific functions of Shisa7 S405 may contribute differentially and/or in a context-specific manner. Although it has been reported that Shisa7 is highly expressed in hippocampal pyramidal neurons, interneurons within the hippocampus also show Shisa7 expression, but the expression patterns differ based on the interneuron subtype (Paul et al., 2017; Zeisel et al., 2015). We also cannot discount the notion that these behavioral alterations are due to compensatory changes in circuits due to global loss of Shisa7 S405 phosphorylation throughout the development. The use of region- or cell-type-specific manipulation of Shisa7 S405 phosphorylation will be valuable in order to fully understand the contributions of Shisa7 S405 to behavior in a circuit-specific context.

In summary, we have expanded on our previous studies further elucidating the importance of the Shisa7-dependent trafficking of synaptic and extrasynaptic GABA_A_Rs in the regulation of GABAergic transmission. Phosphorylation of S405 is not only crucial for the ability of Shisa7 to traffic extrasynaptic α5-GABA_A_Rs, but also for synaptic α2-GABA_A_Rs. Critically, we have discovered that NMDAR-dependent iLTP in hippocampal neurons requires Shisa7 S405 phosphorylation. Moreover, we find that our Shisa7 S405A KI mouse model recapitulates enhanced grooming and locomotor hyperactivity behavioral endophenotypes, suggesting that impaired Shisa7 phosphorylation-dependent trafficking of GABA_A_Rs could be involved in neurodevelopmental disorders.

## Materials and methods

### Animals

All animal handling was performed in accordance with animal protocols approved by the Institutional Animal Care and Use Committee (IACUC) at NIH/NINDS. All mice were housed and bred in a conventional vivarium with *ad libitum* access to food and water under a 12-h circadian cycle (ZT0 - “light on” at 0600, ZT12 - “lights off” at 1800). Time-pregnant mice at E17.5-18.5 were used for dissociated hippocampal neuronal culture. Mice of both sexes at P16-21 were used to prepare acute hippocampal slices for electrophysiology experiments. Adult male mice (2-3 months old) were used for behavioral experiments.

### Generation of a Shisa7 S405A knock-in mouse line by CRISPR-mediated homologous recombination

Substitution of serine with alanine at codon 405 of the mouse Shisa7 gene was achieved by CRISPR-mediated homologous recombination (HR) directly in C57BL/6J zygotes with a single-strand DNA oligo as the recombination template (IDT, Integrated DNA Technologies, Coralville, IA). A few pairs of guide RNAs (gRNA) for SpCas9 (PAM=NGG) were selected based on their relative positions to target codons and by their rankings using DESKGEN (www.deskgen.com), an online gRNA selection tool. gRNAs were synthesized using T7 in vitro transcription as described (Varshney et al., 2015). To select the best pair of gRNAs that can efficiently direct the intended HR, gRNAs were further tested for their in vitro cleavage activities and for insertions and deletions (indel) mutagenesis efficiencies. For the in vitro cleavage assay, genomic PCR products containing the target sites of selected gRNAs were incubated with SpCas9 protein (New England Biolabs, Ipswich, MA) following the manufacturer’s protocol and analyzed on 2% agarose gel stained with ethidium bromide. gRNAs that failed to cleave target DNA fragments were eliminated and the remaining gRNAs were further tested for their efficiencies to induce indels at target sites by Surveyor nuclease assay in an immortalized mouse embryonic fibroblast (MEF) cell line engineered to carry a tet-inducible Cas9 expression cassette as described (Pilato et al., 2012). Upon confirmation of efficient target cleavage activity in these MEF cells, the selected gRNA pair was mixed with SpCas9 protein (PNA Bio, Thousand Oaks, CA) along with a synthetic single-strand donor DNA oligo template as described above. The single-strand donor DNA oligo templates were designed to asymmetrically span the cleavage site and repair the DNA double strand break introduced by the SpCas9 and gRNA (Richardson et al., 2016). The mixture of gRNAs and purified SpCas9 protein were first incubated at room temperature for 15 minutes to load gRNAs into Cas9 protein forming ribonuclear particles (RNPs). The donor oligos were then added to the RNP preparation and microinjected into zygotes of C57BL/6J background as described (Wang et al., 2013). F0 founder mice were screened by PCR and sequencing. The gRNAs pair used to create the Shisa7 S405A variant were 5’-CATAACACGCCGTGGGTT-3’ and 5’-GGAGCACCTTCTGGGTGA-3’, respectively.

Donor oligo sequences carrying the intended substitution is 5’-AGCAGGTGTTCCTGAGACACCAGGCGCGCTCGAGGCAACGTGAACTCATAGCGGGAAGCTCGGCTACCATCACCCAGAAGGTGCTCCTGGGcCATAACACGCCGTGGGTTGGGTGCTGGTGCCAGGC-3’ (127 mer oligo). Mice carrying the S405A allele were genotyped by Sanger-sequencing of a 527 bp amplified fragment generated by PCR using primers 5’-GAGGCTAGAGGAGGAAGGC-3’, and 5’-GGGCATGGTGATGGTGGGC-3’.

### HEK293T Cell Culture

HEK293T cells (ATCC, Cat# CRL-11268) were maintained with culture media containing 1% penicillin-streptomycin (GIBCO), 10% FBS (GIBCO) in Dulbecco’s Modified Eagle’s Medium (DMEM, GIBCO), in a humidified incubator at 37 °C with 5% CO_2_.

### Dissociated Hippocampal Neuronal Culture

Mice hippocampal neurons were prepared from E17.5-18.5 mice embryos of either sex as previously described (Wu et al., 2021b). In brief, hippocampi were dissected from embryonic brains and digested in Hank’s Balanced Salt Solution (HBSS, GIBCO) containing 20 U/ml papain (Worthington) and 100 U/ml DNase I (Worthington) at 37 °C for 45 min. After centrifugation for 5 min at 800 rpm, the pellet was resuspended in HBSS containing 100 U/ml DNase I, and was fully dissociated by pipetting up and down. Cells were then transferred into HBSS containing trypsin inhibitor (10 mg/ml, Sigma-Aldrich) and BSA (10 mg/ml, Sigma-Aldrich). After centrifugation for 10 min at 800 rpm, cells were resuspended in Neurobasal media (GIBCO) supplemented with 2% B27 (GIBCO) and 2 mM GlutaMAX (GIBCO) and were plated on poly-D-lysine (Sigma-Aldrich)-coated glass coverslips or 6-well plates. Cultures were maintained in Neurobasal media supplemented with 2% B27 and 2 mM GlutaMAX in a humidified incubator at 37 °C with 5% CO_2_. Culture media were changed by half volume once a week. Hippocampal neurons at DIV14 were transfected with pCAGGS-IRES-GFP, pCAGGS-Shisa7-IRES-GFP or pCAGGS-Shisa7S405A-IRES-GFP using NeuroMag reagent.

Electrophysiological recordings or immunostaining were performed 48 h after transfection. All transfection kits were used according to the manufacturer’s instructions.

### Electrophysiology

HEK293T cells were co-transfected with cDNA for human α2β3γ2 GABA_A_R subunits together with either pCAGGS-IRES-GFP, pCAGGS-Shisa7-IRES-GFP or pCAGGS-Shisa7S405A-IRES-GFP plasmids using FuGENE® HD (Promega). All recordings were performed 24 h post-transfection. Coverslips containing HEK293T cells were perfused continuously with an external solution (in mM): 140 NaCl, 5 KCl, 2 CaCl_2_, 1 MgCl_2_, 10 HEPES and 10 glucose (pH 7.3; osmolality 285-290 mOsm). The internal solution contained (in mM): 145 CsCl_2_, 10 HEPES, 10 EGTA, 2 MgCl_2_, 2 CaCl_2_ and 2 Mg-ATP (pH 7.3-7.4; osmolality 305-310 mOsm). Recordings were started 1 min after achieving stable, lifted whole-cell configuration at -40mV. GABA (Tocris) was prepared as 1 M stock in the external solution and diluted to 10 mM GABA. Rapid application of GABA or control external solution was performed using a computer-controlled multi-barrel perfusion system (Automate Scientific).

For recording in dissociated hippocampal cultures, neurons were continuously perfused with the extracellular solution containing (in mM): 140 NaCl, 5 KCl, 2 CaCl_2_, 1 MgCl_2_, 10 HEPES, and 10 glucose (pH 7.3; osmolality 300-310 mOsm). The internal solution contained (in mM): 70 CsMeSO4, 70 CsCl, 8 NaCl, 10 HEPES, 0.3 Na-GTP, 4 Mg-ATP and 0.3 EGTA (pH 7.3; osmolality 285-290 mOsm). Whole-cell currents were evoked by saturating GABA (10 mM) at - 70 mV using the same procedure made in HEK293 cells. Miniature inhibitory postsynaptic currents (mIPSCs) and tonic currents were recorded at -70 mV in the presence of 0.5 µM TTX (Alomone Labs), 20 μM DNQX (Alomone labs) and 50 µM D-APV (Abcam). To measure tonic currents, the GABA_A_R competitive antagonist bicuculline (20 μM, Abcam) was bath-applied after obtaining a stable baseline recording at -70 mV and the difference in baseline holding currents before and during bicuculline application was calculated to be the tonic currents.

For recording in acute brain slices, transverse hippocampal slices (300 µm thickness) were prepared from 16-21 days old mice in chilled high sucrose cutting solution that contained (in mM): 2.5 KCl, 0.5 CaCl_2_, 7 MgCl_2_, 1.25 NaH_2_PO_4_, 25 NaHCO_3_, 7 glucose, 210 sucrose and 1.3 ascorbic acid. The slices were recovered in artificial cerebrospinal fluid (ACSF) containing (in mM): 119 NaCl, 2.5 KCl, 26.2 NaHCO_3_, 1 NaH_2_PO_4_-H_2_O, 11 glucose, 2.5 CaCl_2_ and 1.3 MgSO_4_-7H_2_O (pH 7.3; osmolality 300-310 mOsm) at 33°C for 30 min and then were maintained at room temperature prior to recording. mIPSCs were recorded at -70 mV in the presence of 0.5 µM TTX and 20 μM DNQX. After stable baseline recordings, iLTP was induced by transient exposure to NMDA (3 min, 20 μM). The data were binned into 2-min time bins and then averaged to mean amplitude. The extent of iLTP was defined as the ratio of the mean amplitude recorded 30 min after NMDA application to the amplitude recorded before NMDA application.

### Immunocytochemistry

Transfected HEK293T cells grown on coverslips were incubated with rabbit anti-α2 antibody (1:500, Synaptic Systems) in culture medium for 15 min. Next, they were washed briefly with fresh culture medium and fixed with 4% paraformaldehyde and 4% sucrose in PBS. Cells were subsequently incubated with Alexa 555-conjugated anti-rabbit secondary antibody (1:1000, Thermo Fisher Scientific) for the visualization of surface α2. After surface staining, cells were permeabilized with 0.25% Triton X-100, and then incubated with guinea pig anti-α2 antibody (1:500, Synaptic Systems). Alexa 647-conjugated anti-guinea pig secondary antibody (1:1000, Thermo Fisher Scientific) was used for the visualization of total α2. Cultured hippocampal neurons at DIV15 on coverslips were incubated with rabbit anti-α2 antibody (1:500, Synaptic Systems) or rabbit anti-α5 antibody (1:500, Synaptic Systems) in culture medium for 15 min. Next, they were washed briefly with fresh culture medium and fixed with a solution containing 4% paraformaldehyde and 4% sucrose in PBS. Cultured neurons were subsequently incubated with Alexa 555-conjugated anti-rabbit secondary antibody (1:1000, Thermo Fisher Scientific) for the visualization of α2/α5. Coverslips were washed three times with PBS and mounted with Fluoromount-G. Fluorescence images were acquired on a Zeiss LSM 880 laser scanning confocal microscope with a 63 × 1.4 NA oil immersion objective. For quantification, sets of cells were prepared and stained simultaneously. Compared images were acquired at the same time using identical acquisition settings. The fluorescence intensity was analyzed using ImageJ.

### Behavioral Experiments

All behavioral experiments were conducted during the light cycle (“lights on” ZT0-ZT12; 0600-1800) unless otherwise specified. Prior to all behavioral experiments, male mice were allowed 60-90 minutes to acclimate to the behavioral room following transport from the vivarium. Animals were tracked during behavioral tests using AnyMaze software from an overhead camera system. Behavioral equipment was cleaned between subjects with 75% ethanol and allowed to dry for ∼5 min between each behavioral trial.

### Y-maze

The Y-maze apparatus was constructed of clear acrylic and had three arms (height: 12.5 cm; length: 39 cm; width: 9 cm) 120 degrees apart. Mice were placed at the end of one arm and were allowed to explore the maze freely for 5 min. Alternation behavior was defined as consecutive entry into all three arms without repetition and was expressed as percentage of the total arm entries (Hughes 2004). At the end of the test period, mice were placed back in their home cages and the maze wiped and dried before the next animal was run.

### Novel Object Recognition

To acclimate the mice, each mouse was placed in the empty open field box (41 × 41 × 31 cm) and allowed to explore the open field for 10 min, then returned to its home cage. 24 h later, two randomly selected identical objects were placed in the box. Mice were allowed to explore for 10 min and then returned to the home cage. 24 h after the training, one of the familiar objects was replaced with a completely different object in color, size, and texture. Mice were allowed to explore for 10 min. Time spent interacting with each object was analyzed.

### Open Field Test

Mice were allowed to freely explore an open field arena (41 × 41 × 31 cm) for 30 min and their movements were tracked during the test period. Locomotion was operationally defined as the total distance traveled.

### Grooming Behavior

Mice were allowed to habituate to an empty standard mouse housing cage for 10 min. Behavior was then recorded for a total of 10 min and the total amount of time spent grooming was quantified in seconds.

### Nestlet Shredding Test

Mice were placed into a standard mouse cage with bedding and a single nestlet. Mice were then given 60 min of interaction time with the nestlet. After mice were transferred out of the cage, the nestlet was removed and any bedding was carefully removed from the remaining nestlet. Nestlets were allowed to dry overnight prior to weighing the following day. The percent nestlet shredded was calculated using the following formula: % of Nestlet Shredded = (starting weight [g] - ending weight [g])/(starting weight [g]) x 100.

### Marble Burying Test

MBT was conducted as described previously (Han et al., 2019). Briefly, twenty glass marbles were distributed in a 5 × 4 grid. Mice were allowed to bury for a total of 15 min. Latency to bury the first marble was timed within the first 3 min of the test. At the end of the test, marbles with greater than 50% of their diameter remaining were counted and subtracted from 20 to quantify the total number of marbles buried.

### Forced Swim Test

Plastic cylinders were filled with water to prevent mice from touching the bottom (23-25°C) and subjected to the test for a total duration of 6 min. Time spent either swimming or immobile were divided into 2 min epochs (2 min, 4 min, and 6 min) which were then totaled together for total time spent either swimming or immobile. Swimming or immobile behavior were scored offline from recorded videos of the session. Mobility was defined as any movement other than those required for balance and keeping the head of the mouse above the surface of the water (Cryan et al., 2002). Between each session, the water was changed and the cylinder cleaned.

### Homecage Locomotor Activity

Mice were individually housed in a standard home cage under familiar vivarium conditions for a 2-day acclimation period. Activity was then recorded for 24Lh using PAS System (San Diego Instruments). Locomotion was defined as the number of beam interruptions.

### Piezoelectric Sleep Recording

Sleep-wake activity was recorded using a piezoelectric monitoring system (Signal Solutions) as described previously (Wu et al., 2021b). Prior to piezoelectric recording, 2-3 months old male mice were individually housed in a standard home cage under familiar vivarium conditions for a 2-day acclimation period. Sleep-wake activity was then recorded for 24 h. During the recording, mice were left undisturbed and the piezoelectric signals in 2-s epochs were automatically analyzed by a linear discriminant classifier algorithm and classified as sleep or wake. Total sleep percentages and hourly sleep percentages were calculated using SleepStats Data Explorer (Signal Solutions).

### Statistical Analysis

For all biochemical, cell biological and electrophysiological recordings, at least three independent experiments were performed (independent cultures, transfections or different mice). Statistical analysis was performed in GraphPad Prism 9.0 software. Normality distribution was tested by the Shapiro-Wilk test before carrying out a subsequent statistical test. Direct comparisons between two groups were made using either one-tailed/two-tailed Student’s t-test or Mann-Whitney U test. Multiple comparisons were performed using one-way ANOVA, Kruskal-Wallis test or two-way ANOVA with corrections for multiple comparisons.

## Acknowledgements

We are grateful to all members from the Lu laboratory for critical comments on the manuscript. We thank Daniel Abebe at NIH/NICHD for assistance with behavioral tests.

## Author Contributions

K.W., R.D.S., D.C., and W.L. designed the project, and W.L. supervised the project. K.W. performed imaging, biochemical and electrophysiological experiments. R.D.S. and K.W. performed behavioral assays. D.C. recorded GABA-evoked currents in HEK293T cells and neurons. Q.T. performed neuronal cultures. L.D. generated Shisa7 S405A mutant mice. R.D.S., K.W., D.C., and W.L. wrote the manuscript, and all authors read and commented on the manuscript.

---

**Supplementary Figure 1.**
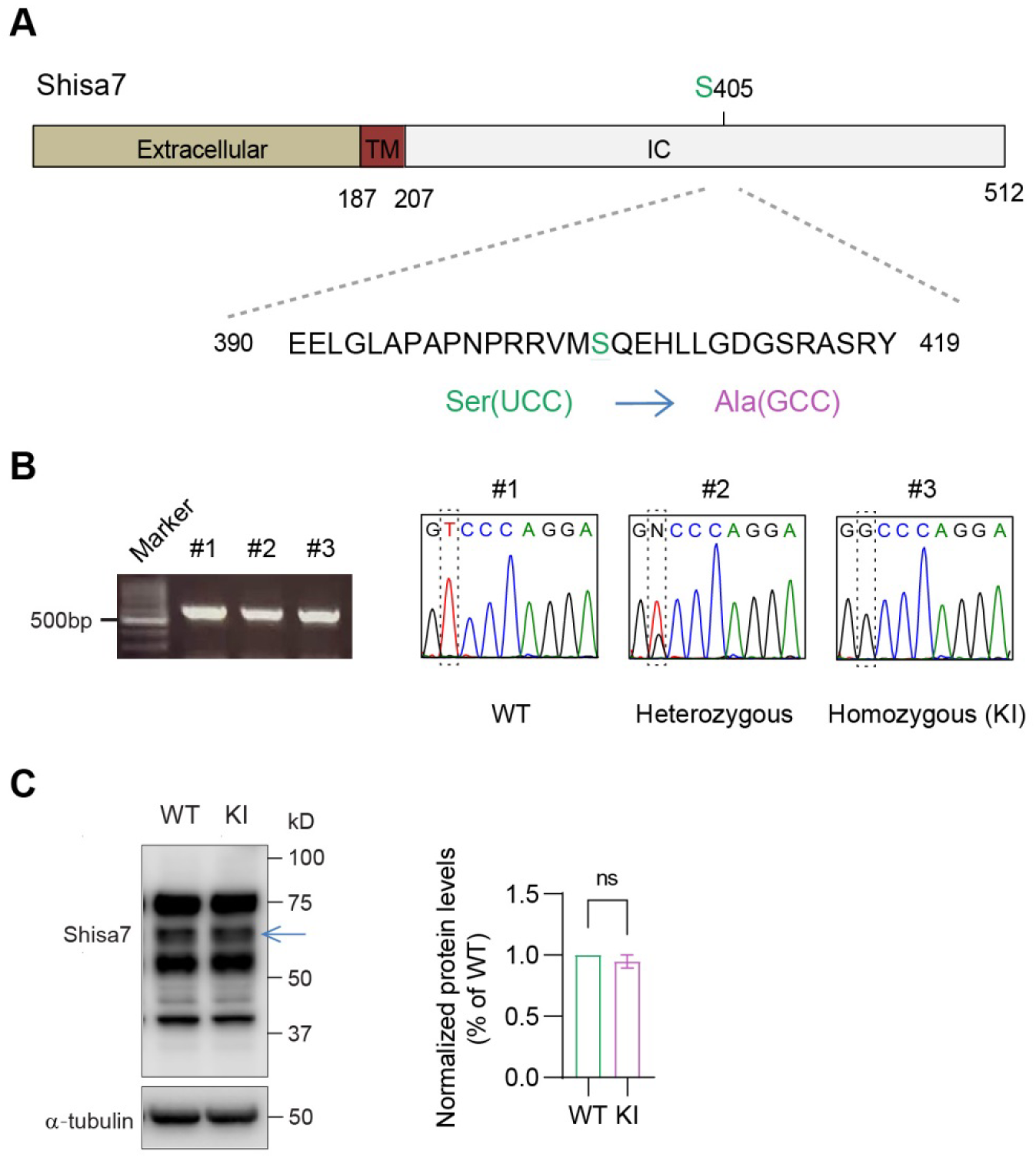
Generation of Shisa7 S405A KI mice. **A**. Schematic of the Shisa7 protein with the WT S405 site in the intracellular region. The serine encoded by UCC (green) in the WT was mutated to an alanine encoded by GCC (purple). **B**. Mice carrying the S405A allele were genotyped by Sanger-sequencing of a 527 bp amplified fragment generated by PCR. As indicated in the dashed box: for WT, only one peak of nucleotide T was observed; for S405A heterozygous, both peaks of nucleotide T and G were overlapped; for S045A homozygous (KI), only one peak of nucleotide G was observed. **C**. Representative western blot of Shisa7 from WT and KI hippocampal lysates using a previously verified antibody raised against Shisa7 (Han et al., 2019). Arrow indicates the band corresponding to Shisa7. There is no difference in Shisa7 expression in total protein lysates from WT and KI mice (n = 4 independent experiments, t test).

